# A comparative genomics and immunoinformatics approach to identify epitope-based peptide vaccine candidates against bovine hemoplasmosis

**DOI:** 10.1101/2020.09.21.305987

**Authors:** Rosa Estela Quiroz-Castañeda, Hugo Aguilar-Díaz, Diana Laura Flores-García, Fernando Martínez-Ocampo, Itzel Amaro-Estrada

## Abstract

*Mycoplasma wenyonii* and ‘*Candidatus* Mycoplasma haemobos’ have been described as major hemoplasmas that infect cattle worldwide. Currently, three bovine hemoplasma genomes are known. The aim of this work was to know the main genomic characteristics and the evolutionary relationships between hemoplasmas, as well as to provide a list of epitopes identified by immunoinformatics that could be used as vaccine candidates against bovine hemoplasmosis. So far, there is not a vaccine to prevent this disease that impact economically in cattle production around the world.

In this work, we used comparative genomics to analyze the genomes of the hemoplasmas so far reported. As a result, we confirm that ‘*Ca*. M haemobos’ INIFAP01 is a divergent species from *M. wenyonii* INIFAP02 and *M. wenyonii* Massachusetts. Although both strains of *M. wenyonii* have genomes with similar characteristics (length, G+C content, tRNAs and position of rRNAs) they have different structures (alignment coverage and identity of 51.58 and 79.37%, respectively).

The correct genomic characterization of bovine hemoplasmas, never studied before, will allow to develop better molecular detection methods, to understand the possible pathogenic mechanisms of these bacteria and to identify epitopes sequences that could be used in the vaccine design.

## Introduction

Hemotrophic mycoplasmas (hemoplasmas) are a group of erythrocytic pathogens of the *Mollicutes* class that infect a wide range of vertebrate animals [1,2]. At first, these small and uncultivable *in vitro* bacteria were classified as species of the genera *Haemobartonella and Eperythrozoon*, within the *Anaplasmataceae* family and *Ricketsiales* order [1]. However, the genetic analysis of 16S ribosomal RNA (rRNA) gene and morphologic similarities showed that these bacteria are closely related to the *Mycoplasma* genus [2,3]. In 2001, the formal proposal was presented to transfer these organisms to genus *Mycoplasma*, within *Mycoplasmataceae* family [4]. Currently, 12 hemoplasma genomes have been identified in the GenBank database, including *Mycoplasma wenyonii* strains and *Candidatus* Mycoplasma haemobos [5–7]. These hemoplasma genomes have provided relevant information about possible pathogenic mechanisms, metabolism and divergences when compared to other *Mycoplasma* species [2]. To date, *M. wenyonii* and *Ca*. M. haemobos have been described as major hemoplasmas that infect cattle worldwide [8–10]. In cattle, acute hemoplasma infections are rare but are characterized by anemia, fever, depression and diarrhea [11,12]. Chronic bovine hemoplasma infections have been associated with variable clinical signs, including low-grade bacteremia, weight loss, decreased milk production, reduced calf birth weight, pyrexia, scrotal and hind limb edema, infertility and reproductive inefficiency, and consequently, the bovine hemoplasmas have caused major economic losses worldwide, mainly when they associate with pathogens of genus *Anaplasma* or *Babesia* [13–15]. In addition, latent and asymptomatic infections have also been reported [14]. Single infections or coinfections between *M. wenyonii* and ‘*Ca*. M. haemobos’ are being reported [16–18]. However, reports of the genomic characterization of bovine hemoplasmas are scarce. So far, two genomes of this species were reported: *M. wenyonii* Massachusetts [5] and *M. wenyonii* INIFAP02 [7] and only one genome of this species was reported: ‘*Ca*. M. haemobos’ INIFAP01 [6]. On the other hand, computational *in silico* tools have been used to design vaccines by rational and cost effective manner, this strategy has several advantages, including prolonged immunity, elimination of unspecific responses and cost-and time-effectiveness [19,20]. In this sense, immunoinformatics is an effective tool that helps to predict and to identify immunogenic sequences and the epitopes that could be recognized by antibodies that induce immune responses [21].

In this work, we analyzed, for the first time, the genomic characteristics and the evolutionary relationships between bovine hemoplasmas. Also, we performed a immunoinformatic analysis to elucidate B-cell epitopes found in several proteins of *M. wenyonii* and *Ca*. M. haemobos that could be used as potential vaccine candidates to prevent bovine hemoplasmosis.

## Materials and methods

### Genome sequences and annotation

The 12 hemoplasma genomes that infect different hosts, included in this study and reported in the GenBank database (https://bit.ly/314fOre), are listed in Table S1. In Mexico, the Anaplasmosis Unit (CENID-SAI, INIFAP) has reported the draft genomes of two Mexican strains of hemoplasmas that infect cattle, ‘*Ca*. Mycoplasma haemobos’ INIFAP01 [6] and *M. wenyonii* INIFAP02 [7]. The general features of 12 hemoplasma genomes were obtained using the QUAST (Quality Assessment Tool for Genome Assemblies) (v5.0.2) program [22] with default settings.

All genomes were annotated automatically to predict the coding sequences (CDS) using the RAST (Rapid Annotation using Subsystem Technology) (v2.0) server (https://bit.ly/2XjTTey) [23] with the Classic RAST algorithm.

The mapping of ribosomal genes (rRNA) was done based on the information reported in NCBI database of genomes of *M. wenyonii* Massachusetts, *M. wenyonii* INIFAP02 and *Ca*. M. haemobos INIFAP01 (NC_018149.1; NZ_QKVO00000000.1; and LWUJ00000000.1, respectively). Transfer (tRNA) RNA genes was carried out using ARAGORN (v1.2.38) (https://bit.ly/3k1R2QT) server [24].The sequence and length of 16S and 23S rRNA genes was obtained from the RNAmmer (v1.2) (https://bit.ly/3glQKCj) server [25].

### Phylogenetic and pan-genomic analysis

For the phylogenetic reconstruction, the 16S rRNA gene sequences of 15 bovine hemoplasmas were aligned with 22 downloaded rRNA gene sequences of other hemoplasma species and two downloaded rRNA gene sequences of the genus *Ureaplasma*, which were obtained from the GenBank database (https://bit.ly/314fOre) using the nucleotide BLAST (Blastn) suite (https://bit.ly/3k2Wkvs) [26]. Multiple alignments between 39 16S rRNA gene sequences were made using the MUSCLE (v3.8.31) program [27]. The jModelTest (v2.1.10) program [28] was used to select the best model of nucleotide substitution with the Akaike information criterion. The phylogenetic tree was estimated under the Maximum-Likelihood method using the PhyML (v3.1) program [29] with 1,000 bootstrap replicates. The phylogenetic tree was visualized and edited using the FigTree (v.1.4.4) program (https://bit.ly/39ROMXV).

Two pan-genomic analyzes were performed using the GET_HOMOLOGUES (v3.3.2) software package [30] with the following options: i) among the 12 hemoplasma genomes; and ii) among the three genomes of bovine hemoplasmas. Briefly, the FAA (Fasta Amino Acid) annotation files of hemoplasma genomes were used as input files by the GET_HOMOLOGUES software package. The get_homologues.pl and compare_clusters.pl Perl scripts were used to compute a consensus pan-genome, which resulting from the clustering of the all-against-all protein BLAST (Blastp) results with the COGtriangles and OMCL algorithms. The pan-genomic analysis was performed using the binary (presence-absence) matrix.

### Comparative genomics

The average nucleotide identity (ANI) values of 12 hemoplasma genomes were calculated using the calculate_ani.py Python script (https://bit.ly/2x96hho) with the BLAST-based ANI (ANIb) algorithm. Ultimately, the level of conserved genomic sequences of bovine hemoplasmas was visualized by alignment the genomes of Mexican strains (‘*Ca*. M. haemobos’ INIFAP01 and *M. wenyonii* INIFAP02) against the reference genome of *M. wenyonii* Massachusetts, using the NUCmer program of MUMmer (v3.0) software package (Kurtz et al., 2004) to get the positions of nucleotides that were aligned; and Circos (v0.69-9) software package [32]. The circular comparative genomic map of bovine hemoplasmas was edited with Adobe Photoshop CC (v14.0 x64) program.

### Prediction of antigenic proteins

After RAST annotation, we identified several proteins of the subsystems, including virulence, disease and defense, cell division and cell cycle, fatty acids, lipids and isoprenoids, regulation and cellular signaling, stress response, and DNA metabolism. To predict antigenicity, the sequence of 11 proteins of *M. wenyonii* and 12 proteins of *Ca*. M. haemobos were submitted to VaxiJen v2.0 server (http://www.ddgpharmfac.net/vaxijen/VaxiJen/VaxiJen.html) with default parameters.

### Prediction of subcellular localization and stability of the proteins

Predicted antigenic proteins of *M. wenyonii* and *Ca*. M. haemobos were submitted to the prediction of secondary structure server Raptor X (http://raptorx.uchicago.edu/).

### Linear B-cell epitope prediction and three-dimensional modelling

B-Cell epitopes were predicted using BCEpred (http://crdd.osdd.net/raghava/bcepred/), and Predicting Antigenic Peptides tool (http://imed.med.ucm.es/Tools/antigenic.pl).

The PHYRE2 server was used to predict the tridimensional structure of the proteins of both hemoplasmas. Phyre2 PDB files were visualized with Protter tool.

## Results

### General features of genomes

The major genomic features of hemoplasmas are shown in Table 1.

**Table 1.** General features of 12 hemoplasma genomes.

| Organism | Assembly level | Length (bp)* | G+C content (%)* | CDS ** | rRNAs<sup>#</sup> | tRNAs<sup>##</sup> |
| --- | --- | --- | --- | --- | --- | --- |
| **‘Ca. M. haemobos’ INIFAP01** | 18 contigs | 935,638 | 30.46 | 1,180 | 3 | 31 |
| ***M. haemocanis* Illinois** | Chromosome | 919,992 | 35.33 | 1,234 | 3 | 31 |
| ***M. haemofelis* Langford 1** | Chromosome | 1,147,259 | 38.85 | 1,595 | 3 | 31 |
| ***M. haemofelis* Ohio2** | Chromosome | 1,155,937 | 38.81 | 1,650 | 3 | 31 |
| **‘Ca. M. haemolamae’ Purdue** | Chromosome | 756,845 | 39.27 | 1,045 | 3 | 33 |
| **‘Ca. M. haemominutum’ Birmingham 1** | Chromosome | 513,880 | 35.52 | 587 | 3 | 32 |
| ***M. ovis* Michigan** | Chromosome | 702,511 | 31.69 | 918 | 4 | 32 |
| ***M. parvum* Indiana** | Chromosome | 564,395 | 26.98 | 578 | 3 | 32 |
| ***M. suis* Illinois** | Chromosome | 742,431 | 31.08 | 914 | 3 | 32 |
| ***M. suis* KI3806** | Chromosome | 709,270 | 31.08 | 856 | 3 | 32 |
| ***M. wenyonii* INIFAP02** | 37 contigs | 596,665 | 33.43 | 678 | 3 | 32 |
| ***M. wenyonii* Massachusetts** | Chromosome | 650,228 | 33.92 | 727 | 3 | 32 |

CDS: coding sequences; \*data obtained with the QUAST program; \*\*data obtained with the RAST server; <sup>#</sup>data obtained with the RNAmmer server; and <sup>##</sup>data obtained with the ARAGORN server.

Of the 12 hemoplasma genomes, ten genomes are assembled in a single chromosome and two draft genomes are assembled in contigs. The genomic features of hemoplasmas allow them to be separated into two groups: the group 1 (previously named *Haemobartonella*) consists of four genomes of ‘*Ca*. M. haemobos’, *M. haemocanis* and *M. haemofelis* species, that have a length from 0.9 to 1.1 Mb, a number of CDS from 1,180 to 1,650, and specifically 31 tRNA genes; and the group 2 (previously named *Eperythrozoon*) consists of eight genomes of ‘*Ca*. M. haemolamae’, ‘*Ca*. M. haemominutum’, *M. ovis, M. parvum, M. suis* and *M. wenyonii* species, that have a length from 0.5 to 0.7 Mb, a number of CDS from 578 to 1,045; and 32 or 33 tRNA genes.

The mapping of rRNA genes shows that hemoplasmas are also separated into two groups. The four genomes of group 1 contain one copy of 16S-23S-5S rRNA operon (Fig 1A). The 16S rRNA gene sequences of group 1 have a length from 1,459 to 1,460 bp. Conversely, seven genomes of group 2 contain one copy of 16S rRNA gene with a length from 1,479 to 1,499 pb, which is separated from one copy of 23S-5S rRNA operon (Fig 1B and 1C). Also, the genome of *M. ovis* Michigan of group 2 contains two copies of 16S rRNA gene with lengths of 1,467 and 3,219 bp, which are separated from each other, and they are separated from the one copy of 23S-5S rRNA operon (Fig 1D).

**Fig 1.**
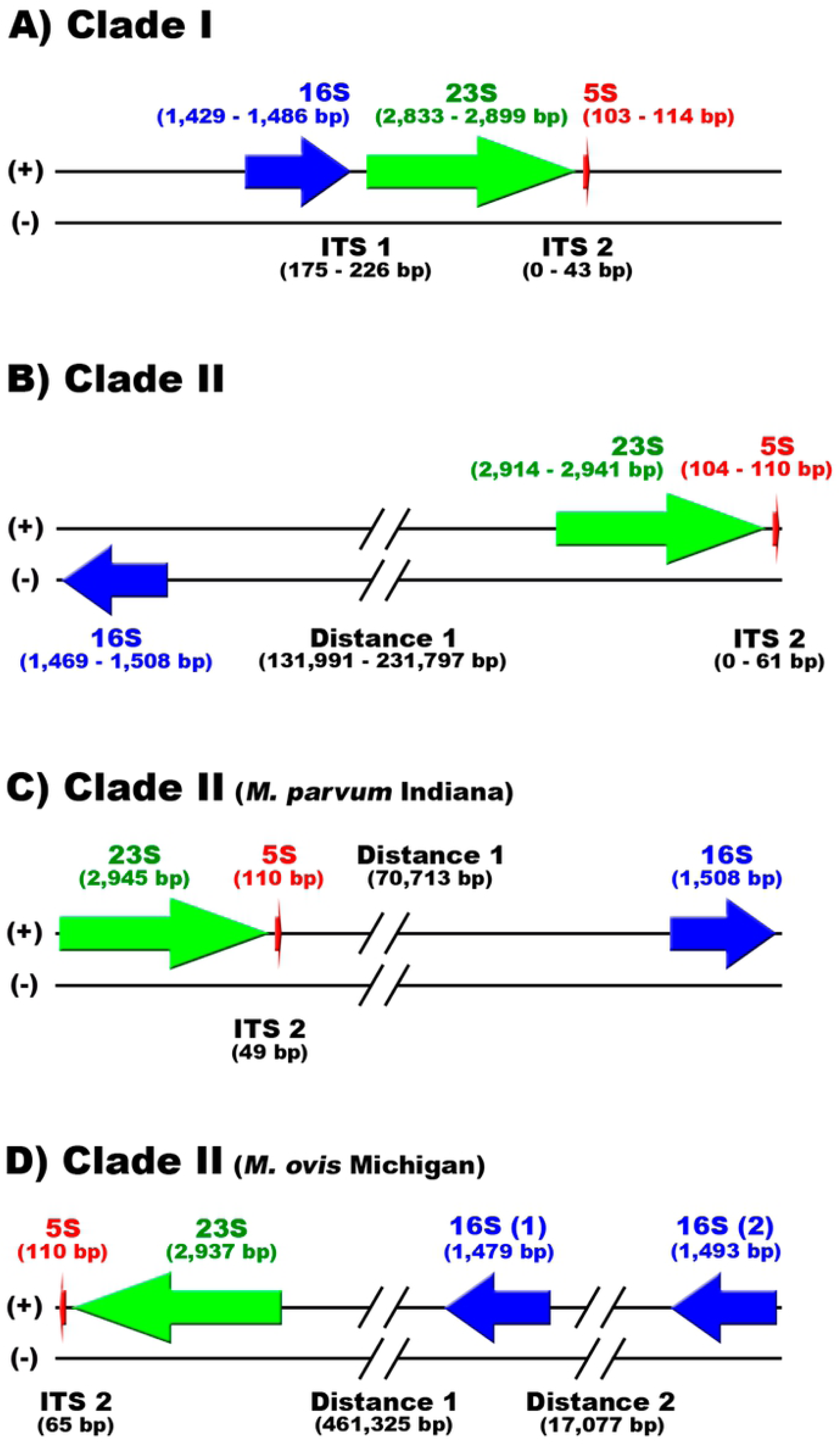
Mapping of rRNA genes of hemoplasmas. A) Genomes of ‘*Ca*. M. haemobos’ INIFAP01, *M. haemocanis* Illinois, *M. haemofelis* Langford 1 and *M. haemofelis* Ohio2 of group 1 contain one copy of 16S-23S-5S rRNA operon. B) Genomes of **‘***Ca*. M. haemolamae**’** Purdue, **‘***Ca*. M. haemominutum**’** Birmingham 1, *M. suis* Illinois, *M. suis* KI3806, *M. wenyonii* INIFAP02 and *M. wenyonii* Massachusetts of group 2 contain one copy of 16S rRNA gene which is separate from one copy of 23S-5S rRNA operon in different chain. C) Genome of *M. parvum* Indiana of group 2 contains one copy of 16S rRNA gene which is separate from one copy of 23S-5S rRNA operon in the same chain. D) Genome of *M. ovis* Michigan of group 2 contains two copies of 16S rRNA gene which are separated from each other, and they are separated from the one copy of 23S-5S rRNA operon in the same chain. The 16S, 23S and 5S rRNA genes are represented by blue, green and red arrows, respectively.

The 16S rRNA gene sequence of ‘*Ca*. M. haemobos’ INIFAP01 has alignment coverage of 82-98% and identity of 98.71-99.93% with ‘*Ca*. M. haemobos’, ‘*Ca*. M. haemobos’ clone 307, ‘*Ca*. M. haemobos’ clone 311 and ‘*Ca*. M. haemobos’ isolate cattle no. 18. Additionally, 16S rRNA gene sequence of ‘*Ca*. M. haemobos’ INIFAP01 has alignment coverage of 99% and identity of 81.83 and 81.73% with *M. wenyonii* INIFAP02 and *M. wenyonii* Massachusetts, respectively. On the other hand, the genomes of *M. wenyonii* INIFAP02 and *M. wenyonii* Massachusetts are very similar in length and G+C content to each other, and they have the same number of tRNA genes and distribution of rRNA genes. However, the 16S rRNA gene sequence of *M. wenyonii* INIFAP02 has alignment coverage of 100% and identity of 97.57% with *M. wenyonii* Massachusetts. Additionally, 16S rRNA gene sequence of *M. wenyonii* INIFAP02 has: i) alignment coverage of 91-98% and identity of 99.24-99.93% with *M. wenyonii* isolate Fengdu, *M. wenyonii* clone 1, *M. wenyonii* isolate ada1 and *M. wenyonii* isolate C124; and ii) alignment coverage of 90-98% and identity of 97.50-97.87% with *M. wenyonii* strain CGXD, *M. wenyonii* isolate B003, *M. wenyonii* isolate C031 and *M. wenyonii* strain Langford.

### Phylogenetic and pan-genomic analyzes

The model of nucleotide substitution of the phylogenetic tree that is based on 16S rRNA gene of hemoplasmas was GTR+I+G. The phylogenetic tree shows that groups 1 and 2 of hemoplasma species are separated into different clades (Fig 2). The clade 1 (blue lines) contains two sub-clades: i) ‘*Ca*. M. haemobos’ species; and ii) *M. haemocanis* and *M. haemofelis* species. The clade 2 (red lines) also contains two subclades: i) ‘*Ca*. M. haemolamae’, ‘*Ca*. M. haemominutum’, *M. ovis* and *M. wenyonii* species; and ii) *M. parvum* and *M. suis* species. The phylogenetic tree topology shows a divergence between two groups of hemoplasmas.

**Fig 2.**
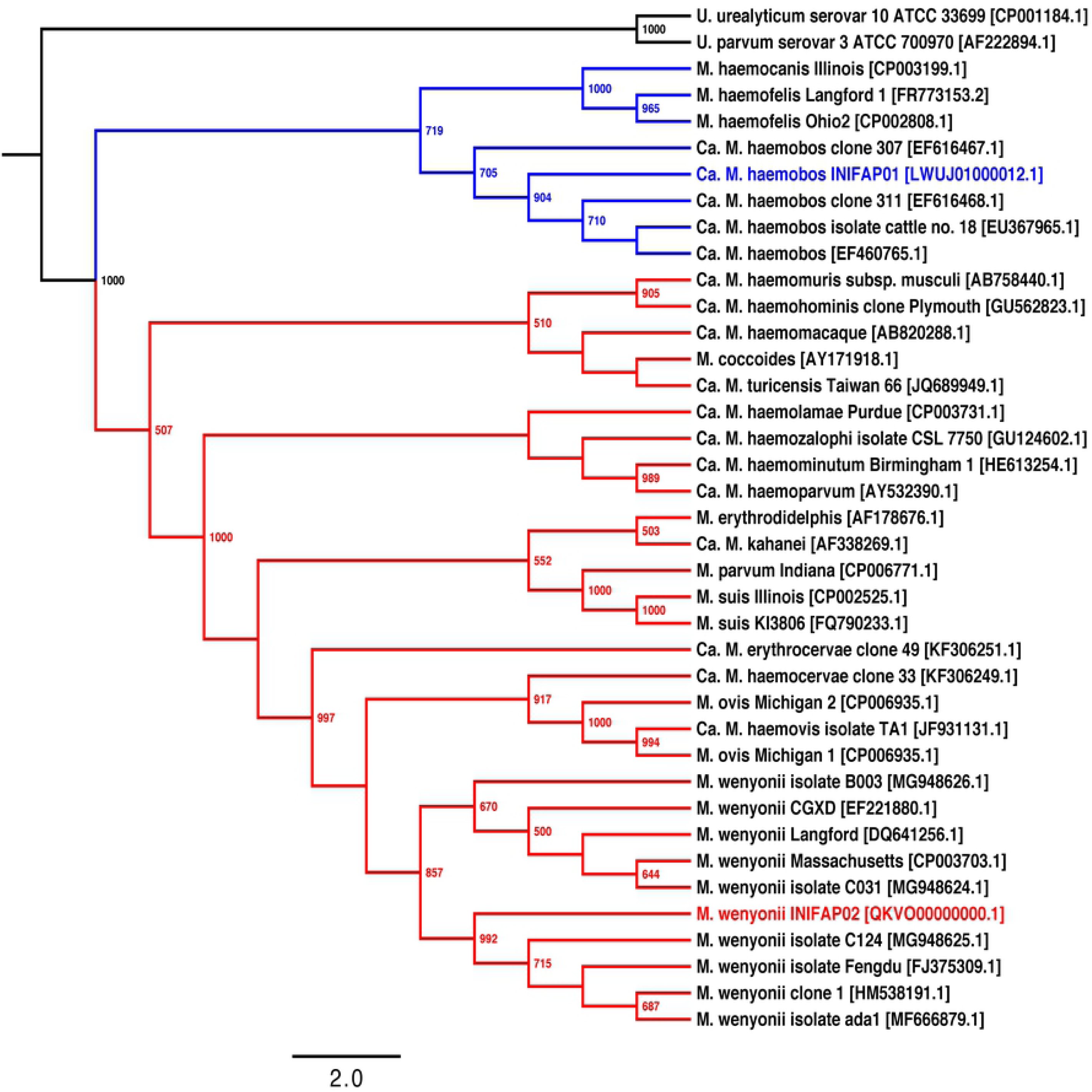
Phylogenetic relationships based on the 16S rRNA genes of group 1 (blue lines) and 2 (red lines) of hemoplasma species. The phylogenetic tree was obtained using the PhyML program with the Maximum-Likelihood method and 1,000 bootstrap replicates. Bootstrap values (>50%) are displayed in the nodes. The model of nucleotide substitution was GTR+I+G. The INIFAP01 and INIFAP02 Mexican strains of bovine hemoplasmas are shown in blue and red letters, respectively. GenBank accession numbers are shown in square brackets.

Pan-genomic analysis among the 12 hemoplasmas shows that the core, soft core, shell and cloud genomes are composed of 110, 146, 787 and 3,099 gene clusters, respectively (Figs S1 and S2). Additionally, the core genomes of groups 1 and 2 of hemoplasmas are composed of 236 and 149 gene clusters, respectively.

Pan-genomic analysis among the three genomes of bovine hemoplasmas shows that the core genome is composed of 154 gene clusters. Also, the two genomes of *M. wenyonii* species share 273 gene clusters. Moreover, ‘*Ca*. M. haemobos’ INIFAP01, *M. wenyonii* INIFAP02 and *M. wenyonii* Massachusetts contain 312, 190 and 157 unique gene clusters, respectively.

### Comparative genomics

ANIb values between different hemoplasma species show that alignment coverage is less than 79% (Fig 3A and Table S2); and identity is less than 83% (Fig 3B and Table S3).

**Fig 3.**
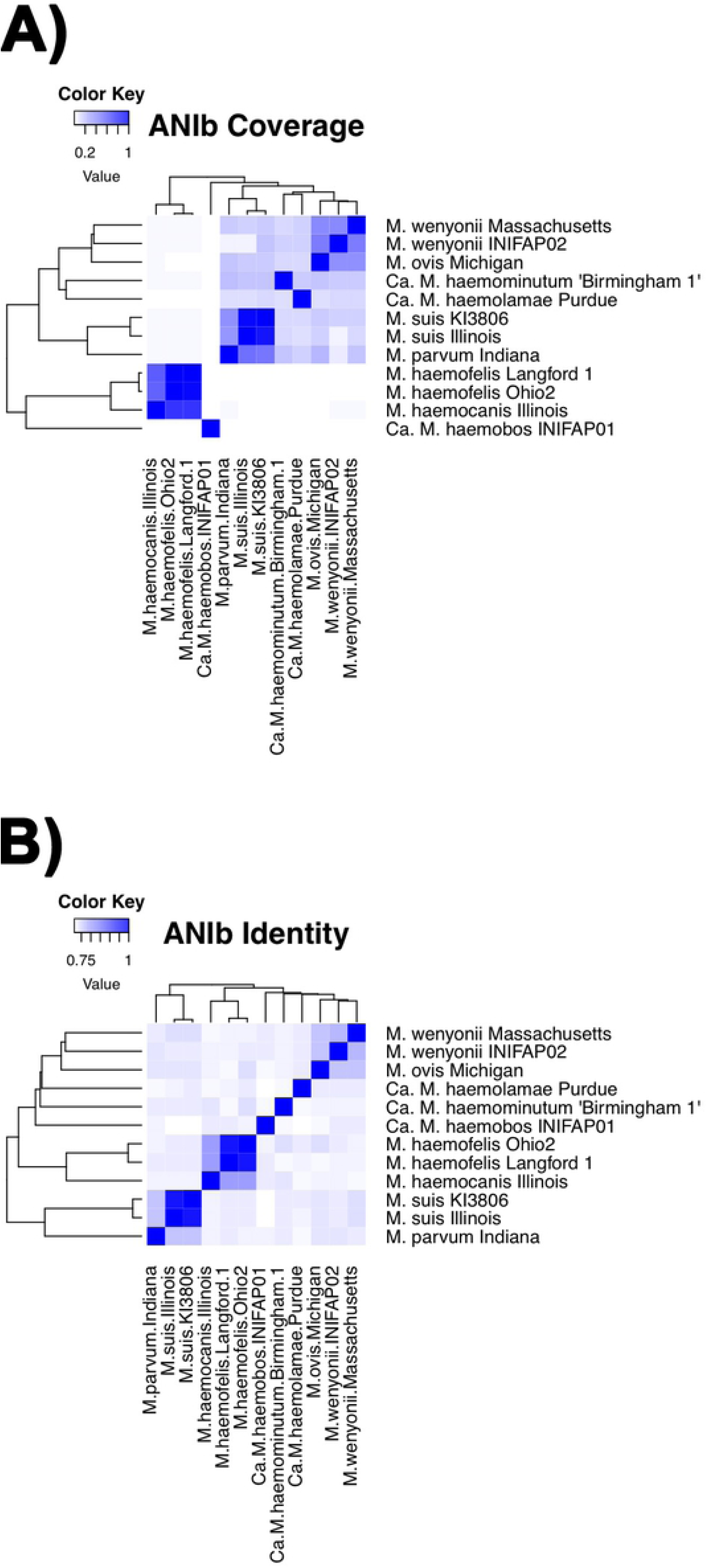
Heatmaps of BLAST-based average nucleotide identity (ANIb) values of 12 hemoplasma genomes. A) Heatmap of ANIb values of alignment coverage. B) Heatmap of ANIb values of identity. Color intensity increases from white to deep blue when ANIb values approach from 0.0 to 1.0 (0 - 100%), respectively.

Also, ANIb values show that ‘*Ca*. M. haemobos’ INIFAP01 has an alignment coverage of 0.46 and 0.31%; and identity of 74.12 and 74.16% with *M. wenyonii* INIFAP02 and *M. wenyonii* Massachusetts, respectively. Moreover, ANI values between the same species show that: i) *M. haemofelis* genomes have an alignment coverage and identity of 97.65 and 97.41%, respectively; ii) *M. suis* genomes have an alignment coverage and identity of 95.13 and 97.63%, respectively; and iii) *M. wenyonii* genomes have an alignment coverage and identity of 51.58 and 79.37%, respectively.

The circular map (Fig 4) shows that ‘*Ca*. M. haemobos’ INIFAP01 genome only has three small regions (red lines highlighted with green marker in inner track) greater than 78% identity that were aligned with *M. wenyonii* Massachusetts genome (black circle in outer track). Also, the circular map shows that *M. wenyonii* INIFAP02 has few regions (blue lines in intermediate track) greater than 78% identity that were aligned with *M. wenyonii* Massachusetts genome.

**Fig 4.**
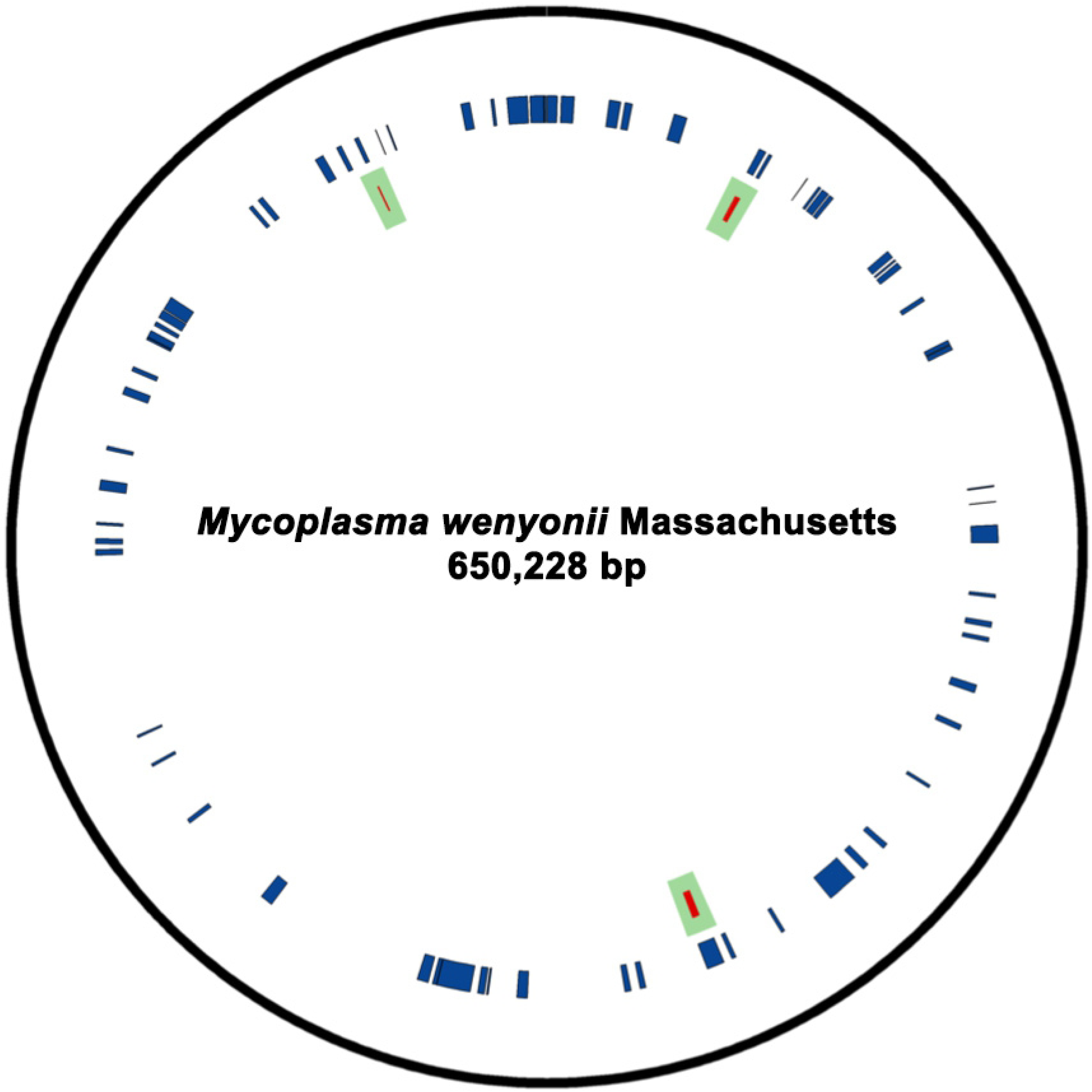
Circular map of *M. wenyonii* Massachusetts and’*Ca*. M. haemobos’ INIFAP01 genomes.

### Selection and prediction of B-cell epitopes in proteins

The sequences of 11 proteins of *M. wenyonii* and 12 proteins of *Ca*. M. haemobos were submitted to VaxiJen. The prediction of Antigen/Non Antigen for each selected protein is shown in Table 2.

**Table 2.**
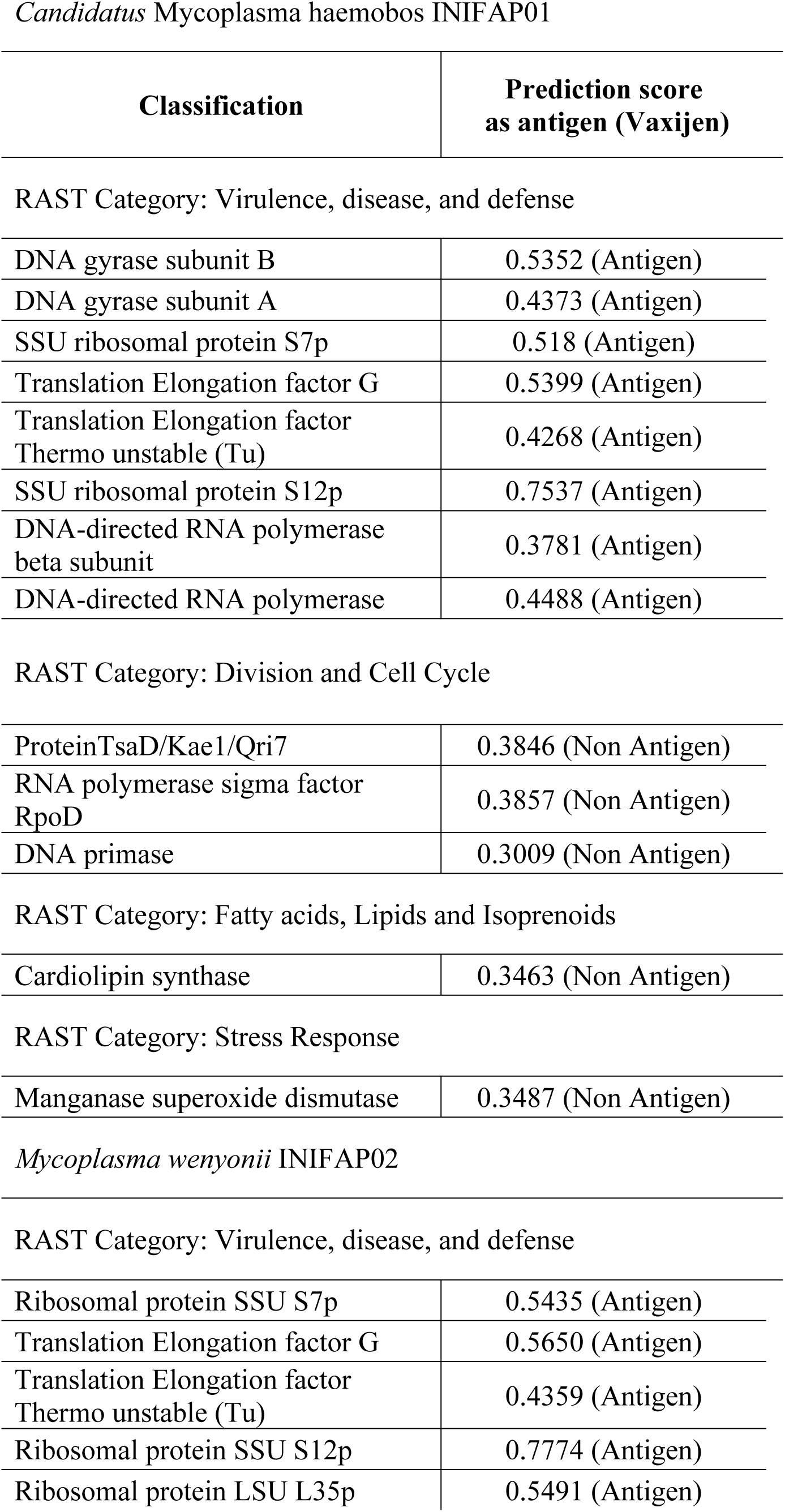
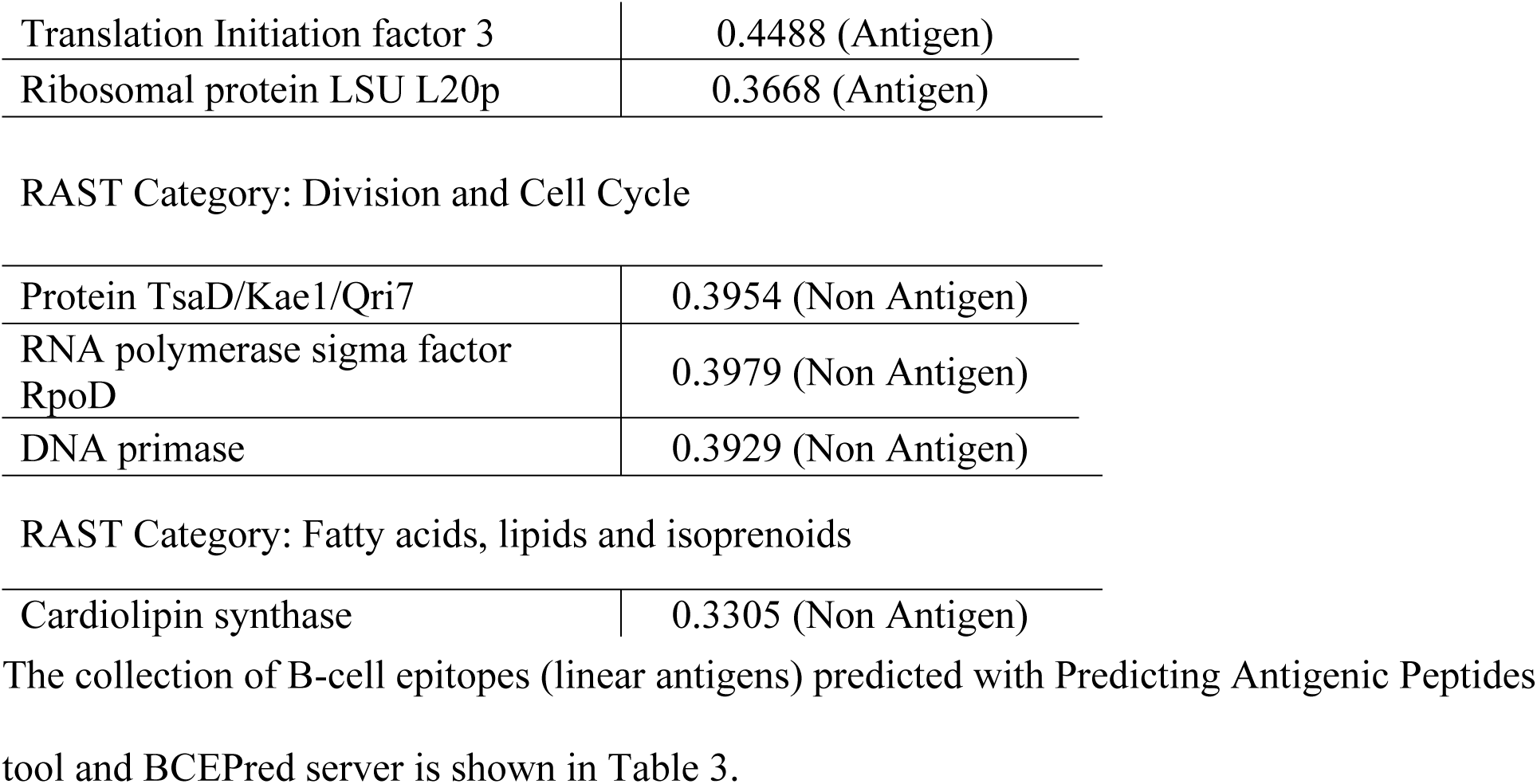
Prediction of antigenicity of proteins of *Ca*. M. haemobos and *M. wenyonii*.

*Candidatus* Mycoplasma haemobos INIFAP01
| Classification | Prediction score<br>as antigen (Vaxijen) |
| --- | --- |
| RAST Category: Virulence, disease, and defense |  |
| DNA gyrase subunit B | 0.5352 (Antigen) |
| DNA gyrase subunit A | 0.4373 (Antigen) |
| SSU ribosomal protein S7p | 0.518 (Antigen) |
| Translation Elongation factor G | 0.5399 (Antigen) |
| Translation Elongation factor<br>Thermo unstable (Tu) | 0.4268 (Antigen) |
| SSU ribosomal protein S12p | 0.7537 (Antigen) |
| DNA-directed RNA polymerase<br>beta subunit | 0.3781 (Antigen) |
| DNA-directed RNA polymerase | 0.4488 (Antigen) |
| RAST Category: Division and Cell Cycle |  |
| ProteinTsaD/Kae1/Qri7 | 0.3846 (Non Antigen) |
| RNA polymerase sigma factor<br>RpoD | 0.3857 (Non Antigen) |
| DNA primase | 0.3009 (Non Antigen) |
| RAST Category: Fatty acids, Lipids and Isoprenoids |  |
| Cardiolipin synthase | 0.3463 (Non Antigen) |
| RAST Category: Stress Response |  |
| Manganase superoxide dismutase | 0.3487 (Non Antigen) |
| Ribosomal protein SSU S7p | 0.5435 (Antigen) |
| Translation Elongation factor G | 0.5650 (Antigen) |
| Translation Elongation factor<br>Thermo unstable (Tu) | 0.4359 (Antigen) |
| Ribosomal protein SSU S12p | 0.7774 (Antigen) |
| Ribosomal protein LSU L35p | 0.5491 (Antigen) |
| Translation Initiation factor 3 | 0.4488 (Antigen) |
| Ribosomal protein LSU L20p | 0.3668 (Antigen) |
| RAST Category: Division and Cell Cycle |  |
| Protein TsaD/Kae1/Qri7 | 0.3954 (Non Antigen) |
| RNA polymerase sigma factor RpoD | 0.3979 (Non Antigen) |
| DNA primase | 0.3929 (Non Antigen) |
| RAST Category: Fatty acids, lipids and isoprenoids |  |
| Cardiolipin synthase | 0.3305 (Non Antigen) |

## Discussion

The hemoplasmas have underwent phylogenetic reclassification after several studies based on molecular markers [33]. Their genome size variation, positional shuffling of genes and poorly conserved gene synteny are evidence of the high dynamic of their genomes [2]. In this work, we found that 12 genomes of hemoplasmas are classified in two groups, and have a different number of CDS and number of tRNAs, additionally, the G+C content vary from 30.46 to 39.27% between the members of the two groups. Thus, G+C content is not specific to each group. Specifically, in bovine hemoplasmas we found differences in genomic features between species. The genome of ‘*Ca*. M. haemobos’ INIFAP01 is significantly longer in length than two genomes of *M. wenyonii* species, but ‘*Ca*. M. haemobos’ INIFAP01 has a lower G+C content. Also, the number of tRNA genes and distribution of rRNA genes are specific to each species. The phylogenetic tree shows that ‘*Ca*. M. haemobos’ INIFAP01 is phylogenetically distant through evolution with *M. wenyonii* INIFAP02 and *M. wenyonii* Massachusetts. Also, the INIFAP02 and Massachusetts strains are closely related through the evolution of group 2.

The number of genes in the core, soft, shell and cloud genomes revealed in the pan-genomic analysis suggest that there is considerable loss/gain of genes through evolution of the 12 hemoplasmas genomes. Also, the genomes of group 1 (four genomes of three species) are more conserved than group 2 (eight genomes of six species); however, to confirm the previous result it is necessary to use a greater number of genomes of different species of group 1. In regard to pan-genomic analysis of bovine hemoplasmas, due to the low number of gene clusters in the core genome it confirms that ‘*Ca*. M. haemobos’ INIFAP01 is a divergent species from *M. wenyonii* INIFAP02 and *M. wenyonii* Massachusetts. In comparative genomics of bovine hemoplasmas, the low percentages of alignment coverage and identity between *Ca*. M. haemobos, *M. wenyonii* INIFAP02 and *M. wenyonii* Massachusetts in the ANI values suggest that *Ca*. M. haemobos INIFAP01 genome has a different structure than genomes of *M. wenyonii* INIFAP02 and *M. wenyonii* Massachusetts.

Surprisingly, the alignment coverage and identity percentages among both genomes of *M. wenyonii* (strains INIFAP02 and Massachusetts) suggest that these strains may not belong to the same species because the ANI values were <95%, the species ANI cutoff value [34–36]. A visual evaluation of circular map suggests that genomes of three bovine hemoplasmas are not conserved between them, in fact, this data confirms that ‘*Ca*. M. haemobos’ INIFAP01 genome has a highly different structure than genomes of *M. wenyonii* INIFAP02 and *M. wenyonii* Massachusetts.

Since bovine hemoplasmas show significant differences at genomic level and they impact in cattle health causing economic losses, we decided to perform an immune-informatic analysis to identify B-cell epitopes that could be used in the design of potential vaccines to prevent bovine hemoplasmosis. The development of vaccines based on this strategy has been successfully used to prevent some diseases in human and animals, including the cytoplasmic protein subolesin used to prevent infestations of tick *Rhipicephalus microplus* [37–40]. The immune-informatics analysis predicts several B-cell epitopes (Table 3), which could be used in the design of molecular detection methods and vaccines. Peptides that contains epitopes have been applied successfully for pathogen detection by serological methods [41,42], immunolocalization of pathogen proteins [43] and vaccines against animal diseases [44–46]. The epitope collection generated in this work will help to design molecular tools that contribute to prevent or diagnose bovine hemoplasmosis.

**Table 3.**
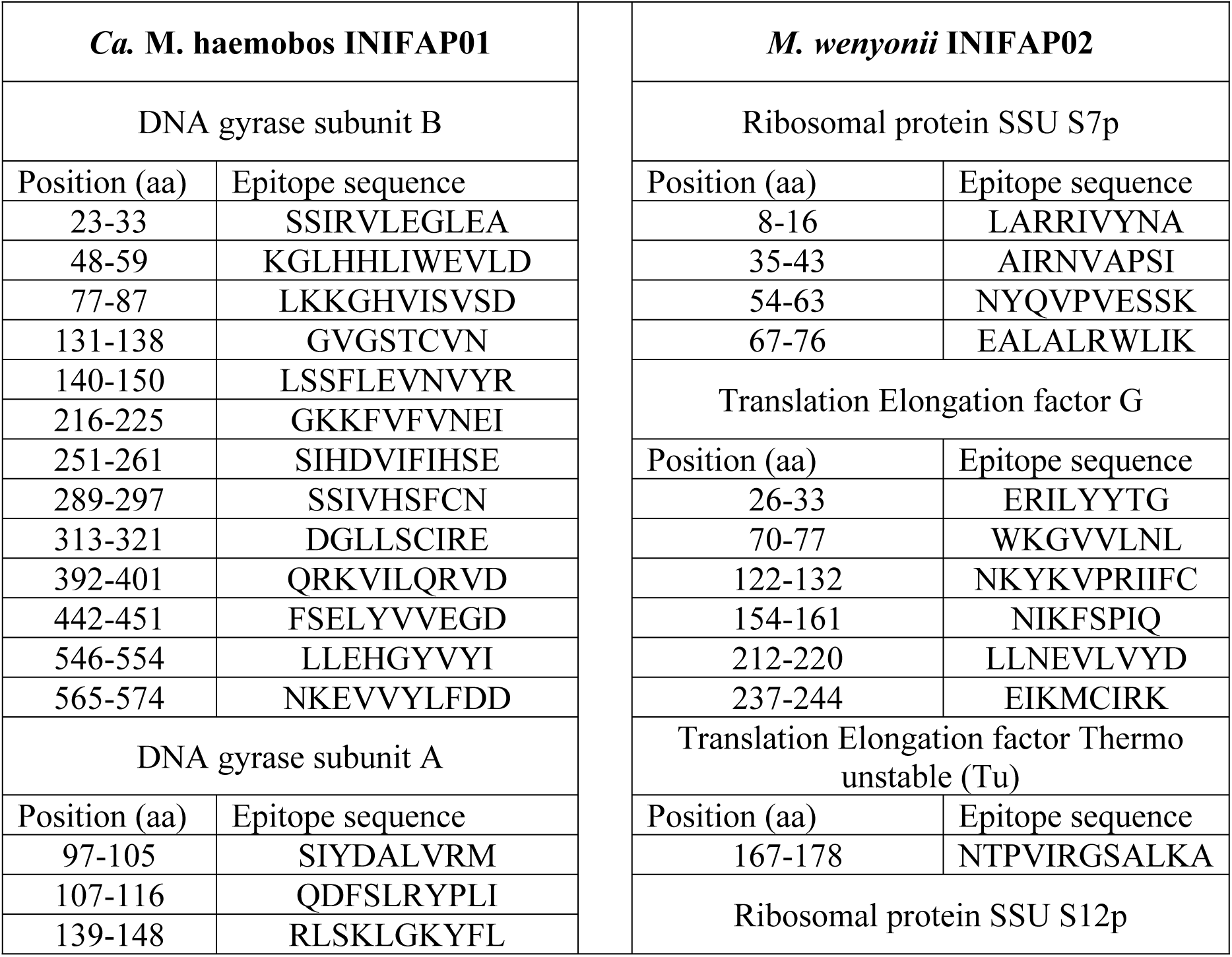
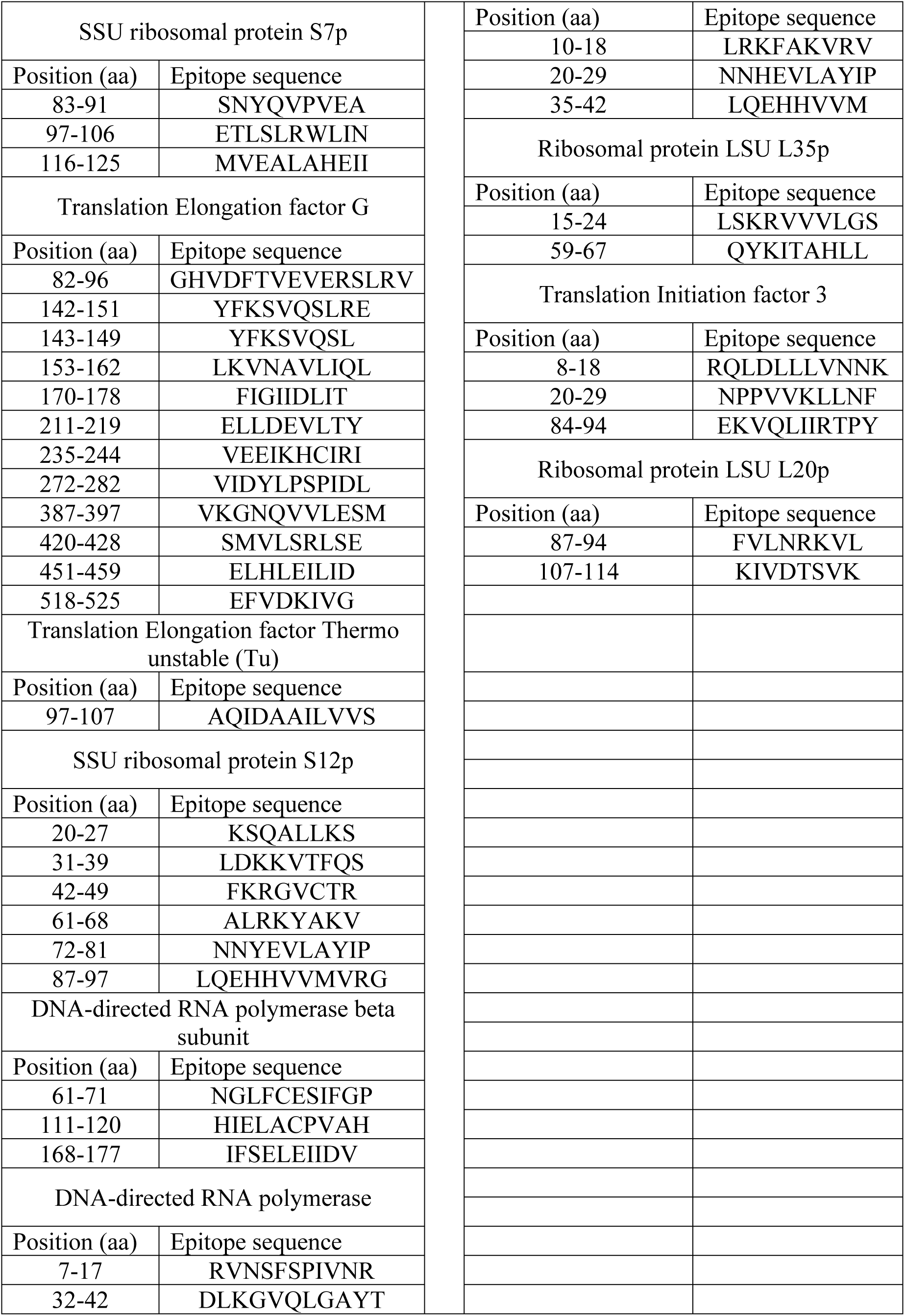
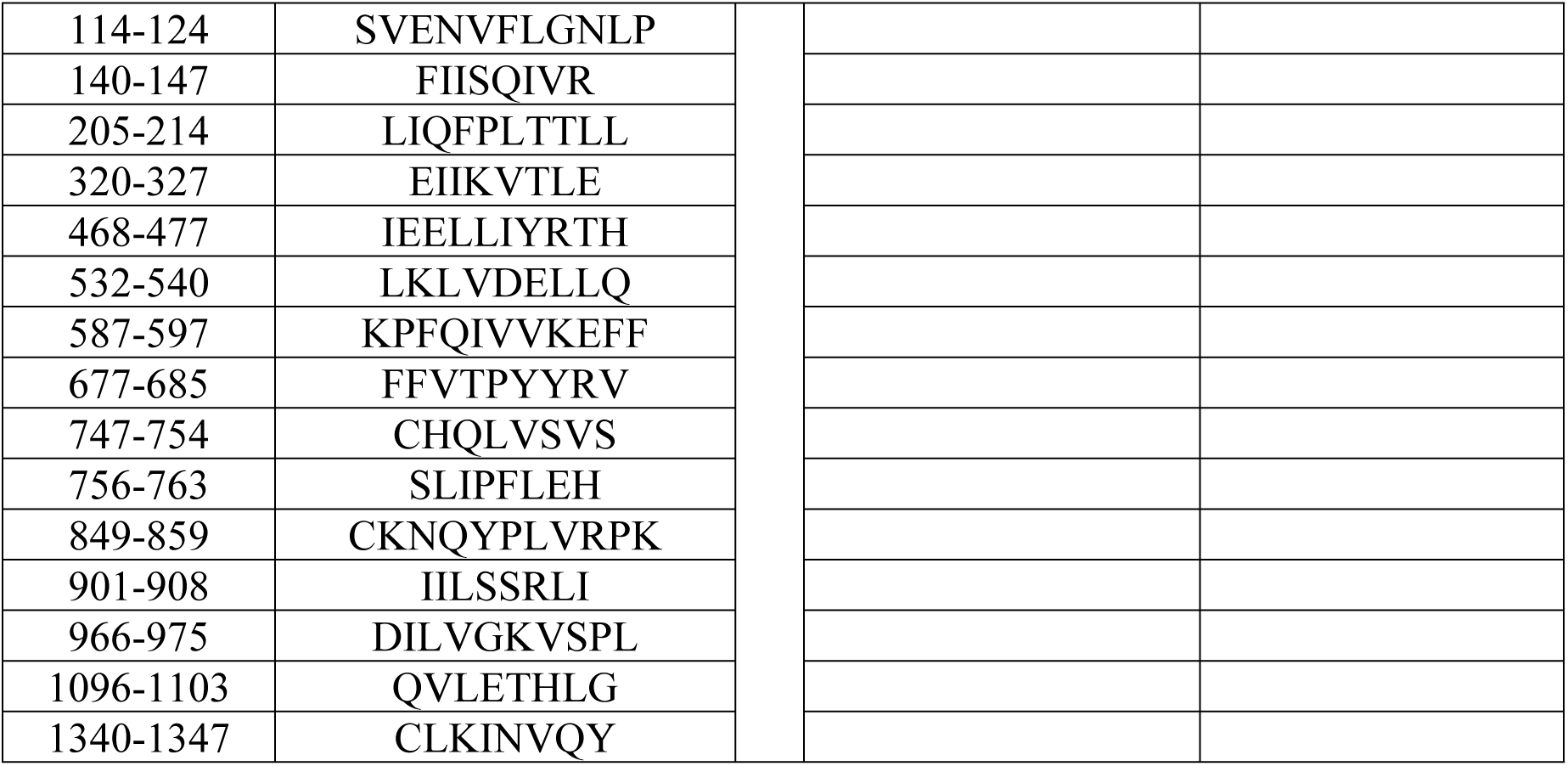
B-Cell epitopes of *Ca*. M. haemobos and *M. wenyonii* predicted by immunoinformatics

## Conclusions

In this work, we described the main genomic characteristics and the evolutionary relationships between three bovine hemoplasmas. This is the first study to report the genomic characteristics of bovine hemoplasma species. Also, the data presented here about antigenic peptides of *M. wenyonii* INIFAP02 and *Ca*. M. haemobos INIFAP01 identified by immune-informatics have potential uses to detect and/or prevent hemoplasmosis.

## Acknowledgements

To Maria Gabriela Guerrero Ruiz (Bioinformatics Analysis Unit, CCG, UNAM) for kindly collaborating with the comparative genomics analysis when using the CIRCOS program. This work was partially granted by Consejo Nacional de Ciencia y Tecnología (CONACyT) Project PN-CONACyT 248855 and CONACyT scholarship 293552.

## Supporting information

**S1_Fig. Pan-genomic analysis of 12 hemoplasmas**.

**S2_Fig. Core, soft core, shell and cloud genomes of hemoplasmas. S1_Table. 12 hemoplasma genomes reported in the GenBank database**.

**S2_Table. BLAST-based average nucleotide identity (ANIb) values of alignment coverage of 12 hemoplasma genomes**.

**S3_Table. BLAST-based average nucleotide identity (ANIb) values of identity of 12 hemoplasma genomes**.

